# Boosting diabetes and pre-diabetes screening in rural Ghana

**DOI:** 10.1101/278960

**Authors:** Bernard Effah Nyarko, Rosemary Serwah Amoah, Alessandro Crimi

## Abstract

**Background:** Diabetes is a growing worldwide disease with serious consequences to health and with a high financial burden. Ghana is one of the developing African countries where the prevalence of diabetes is increasing. Moreover, many cases remain undiagnosed, when along with pre-diabetic cases they can be easily detected. Pre-diabetes condition occurs when blood sugar levels are higher than normal but are not high enough to be classified as diabetes, and it is still reversible.

The main objective of this study is to propose a novel method to increase diabetes and pre-diabetes early detection in rural areas. A secondary aim is to look for new related behavioral determinants specific to rural Ghana, by comparing subjects at risk with those already diagnosed as diabetic.

**Methods:** The screening approach was based on tests performed pro-actively by community nurses using glucometers and mobile phone apps. As a pilot for future policies, glycemic tests were carried out on 101 subjects from rural communities in Ghana deemed at risk and unaware of their diabetic/pre-diabetic status. A comparison of dietary and lifestyle habits of the screened people was conducted in regards to a cohort of 103 diabetic patients from the same rural communities.

*Results:* The pilot screening detected 2 diabetic subjects (2% of the cohort) showing WHO diabetic glycemic values, and 20 pre-diabetic subjects (19.8% of the cohort) which showed the effectiveness of the user-friendly approach. The need for further campaigns about alcohol consumption and physical activities has emerged even in rural areas.

*Conclusions:* Policies based on prevention screening as reported in the manuscript have the potential to reduce diabetes incidence, if actions are taken while patients are pre-diabetic, reduce complication related to late diagnosis and indirectly related health-care costs in the country.

## Introduction

Diabetes is one of the fastest growing non-communicable killer diseases in the world, claiming one life every eight seconds and a limb every 30 seconds [1]. Diabetes of all types can lead to complication in many parts of the body and can increase the overall risk of dying prematurely [2–7]. Pre-diabetes condition occurs when blood sugar levels are higher than normal, but are not high enough to be classified as diabetes; this often has no symptoms, and is reversible [3].

According to the latest 2016 data from the World Health Organization (WHO), among the adults living with diabetes melilites, 80% live in low and middle-income countries such as those in the Asia and Eastern Pacific region. The largest number has been reported in China (90 million people [6]), followed by India (61.3 million people) and Bangladesh (8.4 million people) [5]. Complications of diabetes results in increased morbidity, disability, and mortality and have a high economic cost, especially in developing countries [8]. More specifically, the reported prevalence of type 2 diabetes ranges from 1% in rural Uganda to 12% in urban Kenya. While gestational diabetes has been reported within the Sub-Saharan coutries at different levels (e.g. from 0% in Tanzania to 9% in Ethiopia [10]). Lastly, even considering those values an underestimate, it is expected that the reported cases will reach 82 millions by the 2030 [9].

Ghana is challenged with the increasing prevalence of diabetes, similar to other African countries, with a prevalence of 3.6% in adults and 518,000 diagnosed cases within the country [33]. More specifically, The prevalence of diabetes in some parts of Ghana it has been identified as higher than the world average of 6.4% [11,12]. Moreover, the 2015 report of the IDF indicated many other cases probably remain undiagnosed, posing an increased danger of complications for people living with diabetes unaware of the consequences. Previous studies in the country showed that low level of physical activity and obesity were associated with increased risk of diabetes [4]. Additionally, old age and level of education were also associated with increased risk of diabetes [4]. It has also been observed that within Ghana sugary drinks consumption is linked to type 2 diabetes [13]. Community-based health planning and services (CHPS) is a national health program in Ghana adopted in 1999 to reduce geographical barriers to health care access [14]. According to the CHPS policy, relocating nurses directly to communities could outperform an entire sub-district health center. The cost-effectiveness of CHPS for malaria, diarrhea, and pneumonia has been recently reported [15]. However, specific implementations for non-communicable diseases such as diabetes have not yet been investigated within the CHPS policy.

Vulnerable populations such as those in low- and middle-income countries are generally more affected by diabetes related complications [16]. As in several fields of healthcare, mobile health (mHealth) has the potential of reducing for vulnerable populations with diabetes [17]. This can occur either by sending reminders or by increasing access to patient management [18]. Despite the plethora of studies on mHealth and diabetes management [18,19, 20], no study has been carried out in rural Africa with the aim of improving detection of diabetes and pre-diabetes by using mobile technologies and community nurses.

This study proposes a novel screening approach based on community nurses using glucometers and mobile phones, performing tests on undiagnosed and diagnosed subjects proactively within the community without waiting for participants to present at the clinic. The main objective is to develop a novel method to increase diabetes and pre-diabetes detection, and to find new behavioral determinants related to those conditions. In particular, the purpose of the mobile app is to simplify the tracking operations of the nurses and to collect the data into a centralized secure server. For this purpose, a pilot project was carried out in rural communities of the Central Region of Ghana to assess the feasibility of the approach. A secondary objective was to look for new behavioral determinants related to the rural Ghanaian populations by carrying out a comparison with a group of subjects diagnosed with diabetes from the same communities. Similar to a project carried out in the same area about improving prenatal care [21], the project utilized community nurses instead of the participants to assure reliable glucose level data collection.

## Methods

### Study design

We performed a community-based cross-sectional study using mixed methods of quantitative and qualitative analysis.

Data were collected by community nurses by using a mobile phone application and sent to a secure database. The inclusion criteria for the participants were that they were members of the study communities, the exclusion criteria included being <18 years and having a prior diagnosis of diabetes of any type. A proportional comparison group with diabetes was also recruited, with the aim of possibly finding common dietary habits with the screened subjects found to be diabetic/pre-diabetics in both groups. By the same token, the inclusion criteria for the subjects in the comparison group were of being already diagnosed with Diabetes type-2 in the past, being part of rural communities, and being older than 18 years, No specific age limits were set, as age is not a primary effect, but an indirect effect related to fat deposition and physical inactivity [23]. However, in the view to propose a screening which avoid particular overtime for the nurses during their outreach, we excluded children and young adult under the age of 18 years and we acknowledge this is a limitation of the study.

Subjects were assessed as “non-diabetic” according to their diabetes status awareness, obtained by the following question: "Has a doctor or another health professional ever diagnosed you have diabetes?" Further variables analyzed were: family history (family member diagnosed with diabetes); pregnancy; history of hypertension; screened glucose level; lifestyle characteristics (going to sleep within 1 hour after dinner and level of physical activities); body mass index (BMI), and diet (consumption per typical week of dishes based on staple or maize/corn, root and tuber-potatoes/cassava, and alcohol). Those variables are further described in the following sections as whether they were assessed by the nurses or self-reported.

#### Data source

The research was conducted in the central region of Ghana, specifically within the Biriwa and Anomabo communities. These two communities are in the Mfantsiman Municipal District and based on the 2010 census, the two towns have a population of about 7,500 and 14,389 respectively. Those communities have been chosen due to them being relatively close to a main road connecting urban centers. The assumption is that the members of those communities are more prone to adopt unhealthy habits (such as smoking and drinking) which are more common in urban centers.

Community nurses from rural clinics were instructed to visit rural communities performing glucose screening, when fasted if possible or alternatively at random. Those tested were subjects known to have diabetes, subjects deemed at risk or those willing to be tested. A total of 204 people were tested in a window period of 6 months (from June to December 2017). This sample size was reached following the minimum sample size for the study given by two populations of n=86 subjects. This minimum sample size was computed by using the GPower software (http://gpower.hhu.de/) for an *a priori* two-tail t-test with alpha = 0.05, effect size 0.5 and power 0.90. The quantified sample size also matches constraints according to logistics related to the community nurses. Indeed, the aim was for the proposed screening to be performed by the nurses in addition to their normal activities without resulting in the need for overtime or compromising the other activities were carrying out. As one community nurse per community was used, in this pilot two nurses were employed.

The nurses were equipped with glucometers and low-cost Android smart phones. They received a short (less than one day) training on the app and were supported on its use during the first week of the project. Data were stored through the mobile phone app and sent to a server to improve management and facilitate eventual longitudinal screening. The developed app was based on the <u>CommcareHQ framework</u>, and comprised a series of guided questions that the nurses completed in addition to the glucose test. Some screen-shots of those questions are depicted in Figure 1. If network was not available at the point of data collection, information could be sent later when the network was available. CommcareHQ is a popular mobile data collection, and it has been used in several projects. For a review on projects using the CommcareHQ platform, the reader is addressed to [30].

**Figure 1.**
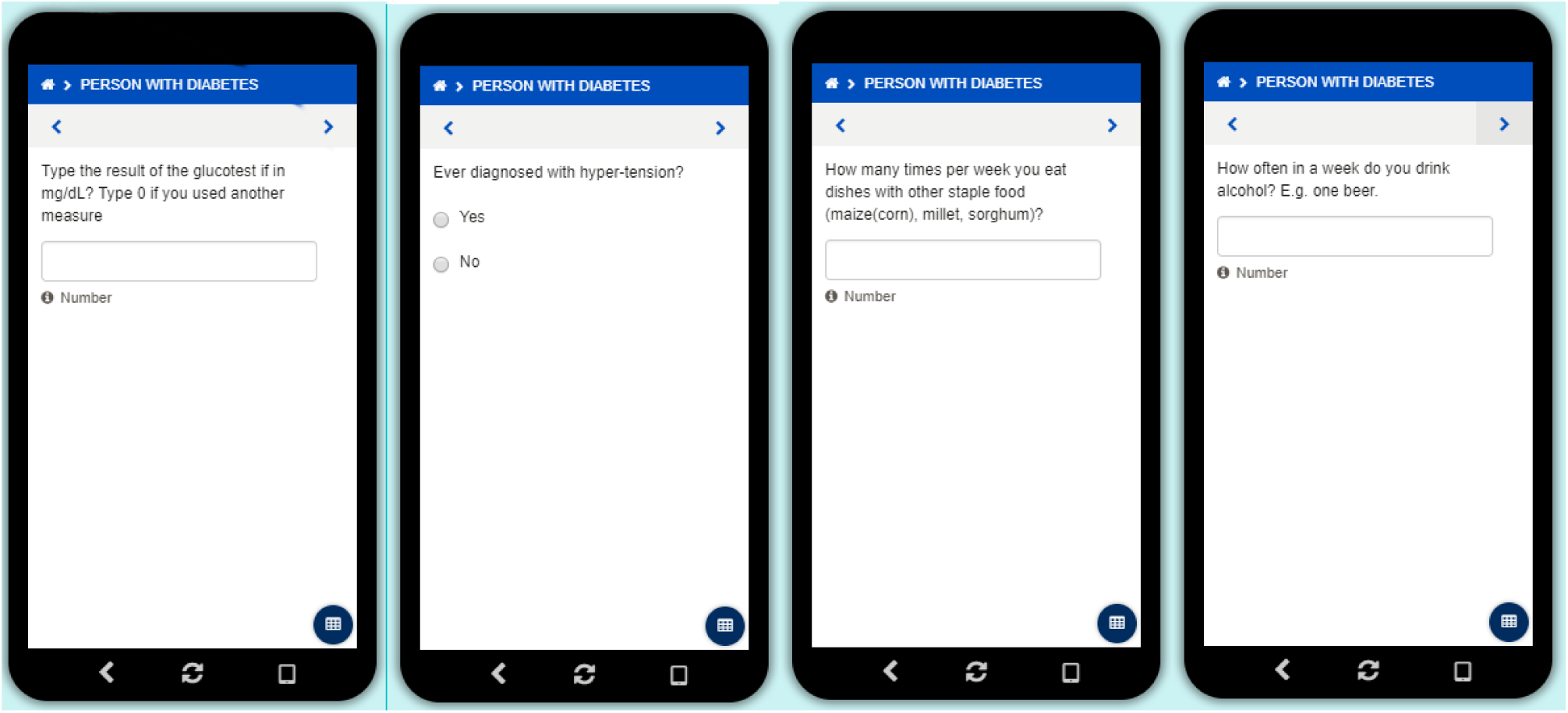
Screen-shots showing some of the guided questions the community nurse were completing while performing the glucose tests. The questions related to diet were referred to the general consuption for a typical week and not the current one.

Figure 2 shows the typical two steps of the screening, first a nurse is performing a glucose test (on the left), and then the data are recorded through the app (on the right). Collected data from respondents were from the rural communities, both male and female, including pregnant women with the intent of capturing eventual gestational diabetes case [22]. The sampling of candidates at risk was based on physical factors and at the nurse’s discretion. During regular visits to the rural communities the nurses assessed whether the person to be tested was a potential diabetic candidate and therefore deemed to be tested (e.g. appearing overweight), paying particular attention to pregnant women. By doing this we have introduced a piloted biases (as overweight people, pregnant women and other cases were deemed at risk by the nurses). Therefore, with those biases, the resulting statistics is not representative of a random screening in the national population.

**Figure 2.**
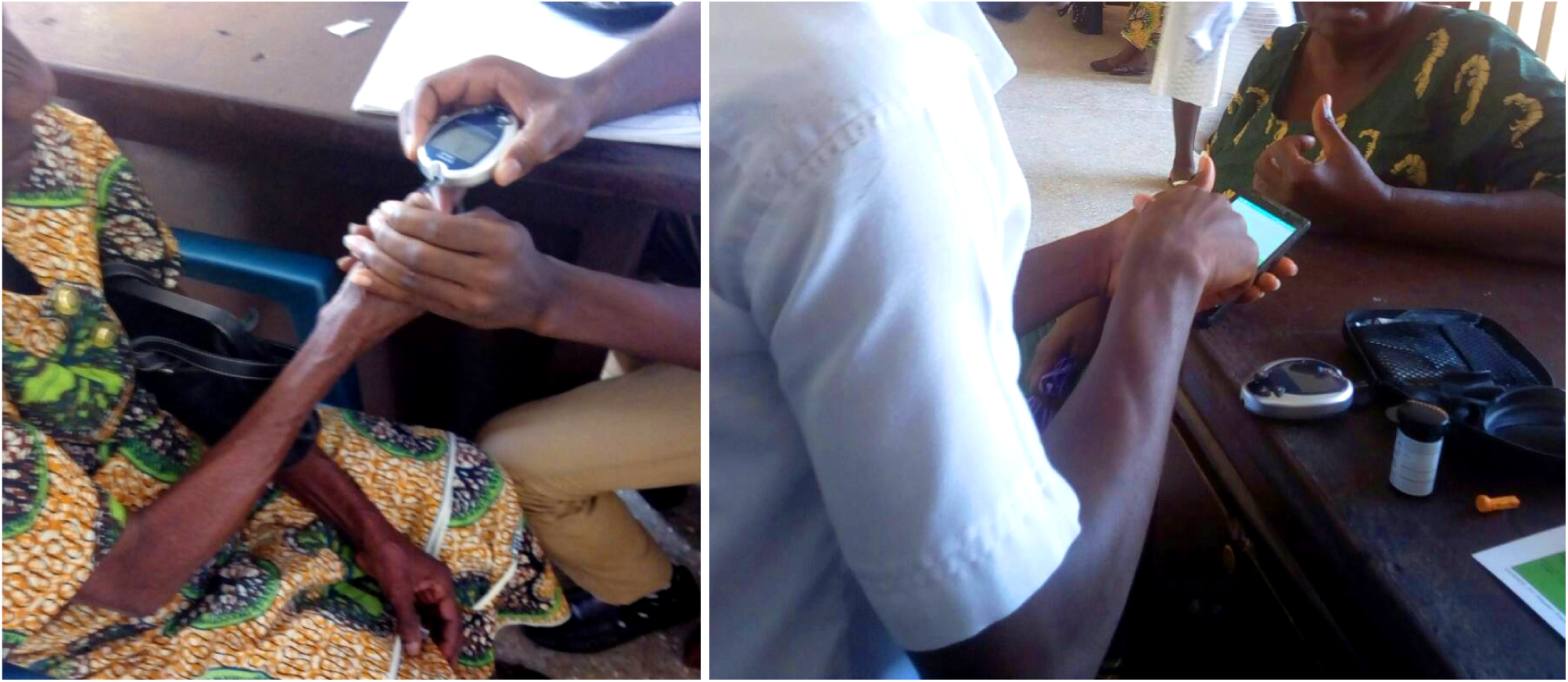
A typical two steps screening, first a nurse is performing a glucose test (on the left), and then the data are recorded through the app (on the right).

Nevertheless, our focus is not to estimate total incidence of undetected diabetes or pre-diabetes, but to propose an approach that can detect as many as possible cases which otherwise would go unnoticed.

However, to avoid a strong bias people deemed healthy that were willing to be tested were included. Any detected diabetic or pre-diabetic cases should be addressed to immediately change their lifestyle or start pharmaceutical therapies. Data were collected on known diabetic patients as well, to evaluate the differences or similarities in lifestyle among the various groups.

With the mobile phone app, the community nurses could also keep track of longitudinal changes or whether a subject has been already tested, in a similar manner of a project conducted in the same area about boosting prenatal care [21]. Subjects who came into contact with community nurses were asked the last time they had a meal, and based on the response blood glucose measurements were classed as either fasting blood sugar levels or random blood sugar levels. Furthermore, the following information were recorded from the subject a) Anthropometric indices (weight, height, BMI) measured by the nurses b) Demographic Information (sex, age) self-reported c) Blood glucose measurement (fasting or random sugar test) measured by the nurses d) Information on risk factors (pregnancy, family history of diabetes, history of hypertension, kidney disorder, alcoholism, low levels of exercise and unhealthy eating habits) self-reported. OneTouch Ultra (Milpitas, CA, USA) and Accu-Check (Hoffmann–La Roche SA, Basel, Switzerland) strips and glucometers were used to measure the blood glucose level. According to the WHO classification [24], a fasting blood glucose level greater 110 mg/dL but less than 126 mg/dL was considered pre-diabetic. A fasting blood sugar level 126 mg/dL or above was considered diabetic. A random blood sugar level greater 140 mg/dL but less 200 mg/dL was considered pre-diabetic. Above 200mg/dL was considered diabetic. Family history of diabetes is referred to occurrence of diabetes in close relatives defined as either father, mother, siblings or offspring. Most of the data were assessed by the nurse though some were self-reported by the subjects and this might represent a limitation. The data entered through the developed Android app were stored in the secure server provided by Dimagi. This cloud-based server is physically located in Cambridge, Massachusetts, USA was used. Data were entered in the forms of the mobile app and transmitted, encrypted, to the cloud-based server, where they were accessed and downloaded via a password-protected web interface.

### Analytical approach

The main aim of the study was to determine if it is possible to easily detect diabetic and pre-diabetic subjects through community nurses already involved and active within the CHPS policy. Additionally, dietary habits and demographic information were collected by the community nurses along with glucose level to explore novel determinants related to the disease, which have not been reported in literature so far.

The data entered through the mobile phone app were downloaded and analyzed by <u>R</u> computing software version 3.5.2. Quality check on the data was performed and forms deemed clearly erroneous were removed. Statistically significant relationships among the collected information were sought comparing the population at risk and the population with assessed diabetes by using a two-tail t-test, and the related p-value less or equal to 0.05 was considered to indicate a significant value.

The study also took advantage of the screening process to collect further insights through qualitative methods. Hence, the quantitative information was complemented with qualitative data obtained through semi-structured interviews performed by the authors with the community nurses to identify further relevant elements at the end of the pilot project. After revision of notes, the transcripts were typed and coded by using <u>the software NVivo</u> 10 (QSR International, Melbourne, Australia). The interviews were thus analyzed by using qualitative conventional content analysis. The starting open questions were “what are your general comments about the projects?”, “Which shortcomings did you notice?”, “What are your suggestions?”.

## Results

The diabetes screening saw over 204 inhabitants in Anomabo and Biriwa over a period of 6 months. Of those, 103 were previously confirmed diabetic participant (CDP) with an average age of 62.9±11.2 years, and 101 people, with an average age of 30 ±9.7 years, with unknown status of diabetes participant (USP). Originally 211 forms were completed, however, 7 of them were deemed erroneous during the quality check and therefore were removed before the analysis. For each person data were collected only once. The CDP cohort comprised 66 female and 37 male subjects, while the USP cohort comprised 95 female and 6 male subjects. Details of some characteristics revealed for the two groups are reported in Table 1. Community nurses see on average 20 patients per day 5 times a week. Their duties mainly encompass malaria, diarrhea, and pneumonia treatments which are generally perceived as more urgent. During the pilot, the total number of people approached for the diabetes screening was 240 with a participation rate of 88%. The 12% of who refused the screening reported as the main motivation the unwillingness to sign the written consent for the study.

**Table 1:** Summary of demographic characteristics for the cohorts reported as mean and standard deviation, (co-) occurrence of hypertension or close relative with diabetes..

| Cohort/Features | Age (year) | BMI | Occurrence of hypertension | Close relatives with diabetes |
| --- | --- | --- | --- | --- |
| confirmed diabetic people (CDP) n=103 | 62.9 ±11.2 | 24.2±5.1 | N=63 (61.2%) | N=72 (69.1%) |
| unknown diabetic status diabetes people (USP) n=101 | 30 ±9.7 | 26.7±4.6 | N = 2 (1.9%) | N = 33 (32.7%) |

Two-sample *t*-test performed comparing the BMIs was statistically significant according to a p-value threshold of 0.05 although they were both normoweight. Some subjects of the CDP cohort also presented with co-occurrence of ulcers (n=4), asthma (n=1), arthritis or rheumatism (n=3), and kidney disease (n=1). At time of testing participants in the USP cohort presented with co-occurrence of asthma (n=4), arthritis and rheumatism (n=1), and typhoid fever (n=1). The subject presenting typhoid fever showed a random glucose blood level of 132 mg/dL which could not be considered neither diabetic nor pre-diabetic. Therefore, typhoid fever was not considered a confounding factor as the tested participant did not show a high value due to this. and the subject was included in the statistics. These results are summarized in Table 2.

**Table 2:** Co-occurrence of disease within the two cohorts.

| Disease | confirmed diabetic people (CDP)<br>n=103 | unknown status of diabetes people<br>(USP) n=101 |
| --- | --- | --- |
| Ulcers | 0 | 0 |
| Asthma | 1 | 4 |
| arthritis or rheumatism | 3 | 1 |
| kidney disease | 1 | 0 |
| Typhoid fever | 0 | 1 |

### Cases detection

During the proactive screening performed by the community nurses, two subjects (1 female, not pregnant, with hypertension, 35.4 BMI; 1 male 25.7 BMI) were found to be hyperglycemic at fasting which would diagnose them as diabetic according to the current WHO threshold [24]. These subjects were not aware of their condition despite close relatives with diagnosed diabetes (son in one case and siblings in the other). They did not present any further symptoms, and they were not habitual consumer of alcohol or red meat. However, their diet was based on dishes with large amounts of maize corn, cassava and rice.

In total, 20 pre-diabetic cases were identified according to the WHO threshold, four tested at fasting and 16 at random (19 female; 1 male). No hypertension or other symptoms were identified, and 5 of had relatives with diabetes (mother or father). Half of the detected subjects reported consuming red meat almost daily, and all claimed to avoid alcohol consumption. They also reported consuming very frequently dishes with maize corn, cassava and rice, averaging respectively 8.5, 8.5 and 3.5 times per week.

All subjects of both groups claimed they were used to perform physical activities due to their daily job. No statistical difference (p-value threshold 0.05) across the two cohorts was detected regarding weekly consumption of red meat, maize corn, cassava or rice. The mean consumption of alcohol across the two populations was also not significantly different (alpha-threshold 0.05), however, in the CDP cohort 14 subjects declared to consume typically at least 1 alcoholic beverage per week (plus 10 claimed to be former alcohol drinkers before their diagnosis and then changed this behavior) while in the USP cohort only 6 participants declared to consume typically at least 1 alcholic beverage per week. Furthermore, in the CDP cohort 3 subjects claimed to have reduced the consumption of cassava based products, and another 2 for red meat. 88 subjects reported to have drastically reduced the consumption of sugar, salt or both but not to have altered their diet. No case of gestational diabetes was detected. Table 3 reports the mean and standard deviation of the blood glucose level for both cohorts, distinguishing whether tests were fasting or random. Summarizing qualitatively, no specific novel determinant was found.

**Table 3.** Mean glucose levels with standard deviation for the two groups and the types of screening and the related p-value.

| Cohort/Glucose level | confirmed diabetic<br>people (CDP) n=103 | unknown status of<br>diabetes people (USP)<br>n=101 | p-value |
| --- | --- | --- | --- |
| Fasting | 160.6 ± 71.6 | 103.1 ± 181 | <0.001 |
| Random | 173.9 ± 65.4 | 123.3 ± 20.6 | 0.002 |

### 3.2. Qualitative comments from the nurses

The two nurses who performed the screening in the rural areas were asked to give their qualitative opinion at the end of the 6 months pilot. The following are extracts of those interviews.

-, Enrolled community nurse, Anomabo Health Centre, Anomabo:

“*There should be continuous education of the masses on diabetes to create awareness. There should also be financial support from governments, and non-governmental organizations to aid routine check up of known diabetes patients, this will encourage them to always take their medication as regular checking of blood glucose level for free help them to know the progress with their condition.*”

“*Medication as insulin is covered by national insurance but not the needles for the screening, and this can make people refrain from screening*.”

-, Enrolled community nurse, Boabab Health Clinic, Biriwa:

“*Most patients were willing to be tested and ready to give any information on their personal lifestyle, diet and medication. There is a general trust in tests performed by clinical personnel regardless on cultural beliefs and on the fact that we are a small clinic*.”

*“Particularly, female clients above the age of forty were pleased to participate in the screening exercise*.”

*“Furthermore, some clients were not able to provide precise information about their diet and family history*.”

“*Screened people reported to perform some kind of sport activity related to their job, believing it was sufficient to keep them healthy*.”

## Discussions

Mobile phone apps and pro-active screening can help community nurses to spot new cases of diabetes and pre-diabetes. In particular, the proposed approach was based on pro-actively performing blood screening during rural visits of the community nurses, who were collecting information via the mobile phone app. This can also help tracking and monitoring, as the nurses can tfollow up the status of the participants in future visits.

Our findings indicate that some individuals in vulnerable populations, such as those in rural Ghana, are not aware of becoming diabetic or being in diabetic condition. We report two cases of diabetic participants in the USP cohort (2%) and 20 as pre-diabetic (19.8%), which are believed to be high numbers compared to previously reported statistcs [34]. However, it must be taken into account that the cohort selection was not purely random, and piloted bias was introduced as the subjects to be tested were chosen according to the nurse’s discretion. Therefore, these percentages should not be taken as a representative sample of the national population. The proposed approach aims instead at detecting as many cases as possible of diabetics and pre-diabetics which otherwise would go unnoticed. For this purpose it proved to be successful, inexpensive and easily integrated into the standard duties of community nurses.

Initially the nurses were equipped with ihealth glucometer dongles for the smartphone (ihealth, Mountain View, CA, USA). The anecdotal comments from nurses were that. despite the initial interest, they were not practical to collect data, though they might be suitable for a single individual. The reason can be related to the familiarity of the nurses to known tools such as Accu-Check and OneTouch, or the cumbersome use of switching continuously between the mobile app of the glucometer dongle and the app to record the data. The motivation behind using a mobile app for tracking data related to diabetes in Ghana is mostly related to the possibility of large institutions to easily monitor status in rural areas, as the data are collectedin a secure centralized server. Moreover, mobile apps appear more user friendly for the nurse in comparison to cumbersome paperwork.

During the qualitative interviews, one nurse pointed out that despite the marginal costs of glucose tests both patients and government are not promoting them, while it could be cost-effective for Ghanaian institutions to detect pre-diabetic cases instead of dealing with a growing diabetic population, as has been shown for similar vulnerable populations [25]. Despite the progresses made in Ghana to achieve universal health access and coverage, financial barriers to diabetes service utilization still exist. Subjects usually refrain from undergoing glucose tests, as they are out-of-pocket.,, and not included in the reimbursements of standard insurances, similarly to Nigeria and Tanzania [26]. The other nurse mentioned that subjects were often not able to report their dietary habits, and they were not aware of the impact they had on their health. Conversely to prenatal care [31], it appears that generally the members of the communities trust the personnel of the clinics for this type of tests.

At population level the CDP and USP cohort were normoweight (having a BMI between 18.5 and 24,99), though there were subjects which were obese and with hypertension in both groups. It is worthwhile to mention that the CDP subjects might have some dietary changes already after being informed of being diabetic, which could have affected their weight and other measurements. However, from the interviews it seems that the major changes were the reduction of consumption of salt and sugar, some increased their consumption of vegetables and fruit, and some reducing the consumption of alcohol beverages. The study did not track behavioral changes, but we can presume that these may have occurred and can represent confounding factor. No statistically significant difference in alcohol consumption between the two groups was detected. However this could be due to the fact that some individuals in the CDP cohort changed their dietary habits (10 people reported to have cut out alcohol consumption after their diagnosis). Moreover, the nurses were instructed to focus for both groups on women which might consume less alcoholic beverages than men. Nevertheless, the general impression of the nurses was that the alcohol consumption had an impact. With the increase in quality of life in the country, western habits such as alcohol consumption might be also increasing, and therefore augmenting the risk of diabetes. It has been noticed that people informed of their condition tend to change their dietary habits. Nevertheless, but given the country-wide growing trend in alcohol consumption, social marketing campaign related to this [27] should be performed.

Almost all subjects of both groups reported to perform regular physical activities related to their job. However, this information seems vague and it is not clear whether this physical activity is aerobic or resistance or whether it is sufficient to keep a healthy glucose level. Most likely further activities should be proposed. Jogging and other sport activities are inexpensive and easy to promote. Therefore, promoting this type of sport activities can address this issue. Moreover, it appears necessary in the future to use more detailed investigations about sport activities, such as using the WHO global physical activity questionnaire [32].

Ghana has experienced an exponential increase of the mobile network, social media and smartphones in the recent years [28]. Beyond the screening of the population carried out by nurses, smartphones can have an impact on glycemic control, as smartphone dongles can be inexpensive and attractive to young users. Strategies such as gamification, and social media should be explored to increase awareness on glycemic levels as shown in other contexts [29].

## Conclusions

Proactive glycemic screening on vulnerable population – such as those living in rural areas – can be effective in detecting new cases of diabetes and pre-diabetes. Our approach using community nurses screening subjects deemed at risk and collecting data on mobile phone was found to be effective, and suitable for longitudinal studies. Campaigns increasing awareness of alcohol consumption, physical activity, nutrition and healthy habits should be emphasized in any prevention strategy as the population seems to still be unaware of the consequences.

Despite this, studies with larger population are required to confirm the results. The diabetes and pre-diabetes screening described in this manuscript can be easily included into the national CHPS policy with several potential benefits. Those benefits include reducing incidence by detecting cases of pre-diabetes which hopefully will not convert into type-2 diabetes and enabling timely treatment of diabetes patients avoiding complications related to delays in treatment.

## Ethics and consent

All procedures performed in studies involving human participants were in accordance with the ethical standards of the institutional and/or national research committee and with the 1964 Helsinki declaration and its later amendments or comparable ethical standards. Written consent for the reported data was collected.

## Acknowledgements

This research was conducted with the support of Baobab medical center in Biriwa (Ghana) and Anomabo Health Center (Ghana) and the related communities.

## List of abbreviations

IDF: International Diabetes Federation
WHO: World Health Organization
CHPS: Community-based health planning and services
BMI: body mass index
CDP: confirmed diabetic people
USP: unknown status of diabetes people

